# When Attention Hurts: The Effect Of rTMS On Neural Correlates Of Time Perception

**DOI:** 10.1101/2024.09.27.615494

**Authors:** Federica Contò, Giulia Ellena, Michele Tosi, Nicholas A. Peatfield, Lorella Battelli

**Affiliations:** Center for Neuroscience and Cognitive Systems@UniTn, Istituto Italiano di Tecnologia, Via Bettini 31, 38068 Rovereto (TN), Italy; Centro Interdipartimentale Mente e Cervello, University of Trento, Rovereto, Italy; School of Engineering, Simon Fraser University, Burnaby, BC, Canada; Biomedical Physiology and Kinesiology, Simon Fraser University, Burnaby, BC, Canada; Berenson-Allen Center for Noninvasive Brain Stimulation, Department of Neurology, Beth Israel Deaconess Medical Center, Harvard Medical School. Boston, MA, USA; Cognitive Neuropsychology Laboratory, Department of Psychology, Harvard University, Cambridge, MA, USA

**Keywords:** time perception, attention, transcranial magnetic stimulation

## Abstract

The parietal lobe plays a crucial role in the attentional networks that help shape our perception, including our perception of time. Based on neuropsychological and neurophysiological evidence, a “when” pathway including the right parietal lobe has been proposed as the critical cortical site for the discrimination of objects across time. When an oddball stimulus is presented in a stream of identical standard stimuli, it is perceived as lasting longer, even when its actual duration is shorter. While attentional capture seems to play a major role in this subjective expansion of time the cortical mechanisms responsible for this effect remain unclear. We therefore set to investigate the direct role of parietal brain areas in time perception using combined repetitive transcranial magnetic stimulation (rTMS) and electroencephalogram (EEG). We measured the perceived duration of an oddball stimulus in each participant before and after inhibitory 1-Hz rTMS stimulation at one of three scalp locations: the right intraparietal sulcus (rIPS), the right inferior parietal lobe (rIPL), or the occipital cortex as a control. EEG was recorded throughout. Stimulation over the rIPL caused a more veridical experience of times subjective expansion towards the oddball, while rTMS over the rIPS and the occipital cortex had no effect. These data provide theoretically challenging notions to the concept of the cortexes role within time perception and the mechanisms involved.

## 1. Introduction

The ability to correctly perceive time is an essential component of our conscious experience. However, time perception illusions are common not only in the laboratory [1] but also in real-life, as when time famously slows during traumatic events [2,3]. This subjective experience, alongside extensive experimental investigation, has led to a popular conceptualization of time perception as a meta-percept, dependent on the integration of input from multiple senses rather than determined by any single ‘clock’ in the brain [4,5].

Time perception has been investigated using *oddball* paradigms, where a unique, unpredictable, infrequent stimulus is presented among homogenous standards [6–9]. Results show that the oddball is perceived as lasting substantially longer than the standard even when there is no real time difference[6]. Tse and colleagues argued that the involuntary deployment of attentional resources to the oddball impacts the accumulation of information about this target, ultimately leading to an increase in perceived duration. However, there is a strong alternative to this *attentional capture* hypothesis. The neuronal repetition theory of time perception suggests that perceived time is a function of the required “neuronal energy” to represent a stimulus [10]. A core implication is that the subjective duration of a repeated stimulus will be contracted due to a reduction in elicited neural activity caused by adaptation. According to this *distractor dilation* account, a relative increase in the perceived duration of an oddball reflects a gradual dilation in the perceived duration of standard stimuli rather than any discrete effect on the perception of the target itself.

Here we use combined rTMS and event-related potentials to investigate the nature of time perception and to discriminate between the *attentional capture* and *distractor dilation* theories of the oddball temporal illusion. In each of two experiments, we had participants judge the duration of visual oddballs and standards following 1-Hz rTMS of parietal cortex. Recent results suggest a key involvement of inferior parietal lobe (IPL) in both time perception and the deployment of attention in time, but there is still no evidence directly linking the oddball illusion to specific sites within the parietal lobe. We used 1-Hz rTMS to deactivate cortical functioning in the posterior and inferior parietal cortex [11], subsequently examining the impact of this manipulation on the oddball temporal illusion. If the oddball illusion reflects attentional capture, our expectation was to see it - and its correlate in the visual ERP - reduced following stimulation. In contrast, if the illusion is a product of sensory adaptation and of the dilation of perceived distractor duration, stimulation of parietal cortex should have no effect.

We approached our electrophysiological data with interest in the P3 complex of the visual ERP [12] and the P3b in particular, a traditional ERP signature of an oddball[13]. This component has a strong tempo-parietal topography and is sensitive not only to attentional modulation, but also to the temporal qualities of eliciting stimuli like regularity[14]. If the oddball temporal illusion is a product of attentional capture, and this creates a difference in the P3b, our expectation was that rTMS deactivation of parietal cortex would reduce the amplitude of this index of capture. The first target of our study was the posterior inferior parietal sulcus (IPS), traditionally considered a main cortical hub of visual attention[15]. However, more inferior portions of the parietal lobe (IPL) have shown to be pivotal in attention-based relative timing[16], the mechanism potentially implicated in the subjective expansion of time phenomenon.

## 2. General procedure

In each of the two experiments described below participants completed two sessions of an oddball time perception task. They viewed an ongoing stream of visual stimuli, most of which were blue (standards) and a few of which were red (oddballs), and reported via button press whether each oddball had longer or shorter duration than the standards. Oddballs occurred in about 10% of trials, and while oddballs had a duration that varied as a function of performance, standards had a fixed duration of 1 second. In each session, participants completed two task runs, one before and one after 10 minutes of 1-Hz rTMS. Each participant completed an experimental session involving stimulation of the parietal lobe (rIPS in Experiment 1; rIPL in Experiment 2) and a session involving stimulation of the occipital pole as a control, counterbalanced across sessions.

## 2: Experiment 1

### 2.1 Methods

#### 2.1.1 Participants

Eighteen healthy subjects (9 women) aged 21-31 years (mean 25) gave informed consent before taking part in the study. All participants were right handed, had normal or corrected to normal vision and had no history of neurological or psychiatric disorders. The study was approved by the ethical committee of the University of Trento in compliance with national legislation and the Declaration of Helsinki.

#### 2.1.2 Stimuli & Apparatus

The task was presented on a 22” Samsung 2233RZ LCD monitor running at 120Hz using EPrime 2.0 in the Windows 7.0 operating system. All stimuli were presented on a black background (0.5 cd/m^2^), the oddball stimuli were red (18.4 cd/m^2^) and the standard was blue (15.2 cd/m^2^). The stimuli had diameter of 6 degrees visual angle and were presented centrally.

#### 2.1.3 Procedure

##### 2.1.3.1 Time Perception (TP) Task

Participants sat 57 cm from the monitor with their head positioned on a chin rest and were asked to maintain their eyes at the center of the computer monitor. A brief instruction screen appeared at the beginning of each run. Each run lasted ∼7 minutes and was split into two blocks separated by a 30 sec break. Every 8-11th stimulus was an oddball and participants reported whether the oddball was of longer or shorter duration than the preceding standard by pressing one of two keys on the keyboard. These keys were z and m on the international keyboard layout, and were counter-balanced between participants. Standard stimuli were presented for 1,050 ± 50 msec with an inter-stimulus-interval of 508 ± 50 msec. The duration of the oddball was controlled by a weighted two-up one-down adaptive staircase procedure designed to measure the 25th and 75th percentile of the psychometric function. The starting duration of the oddball were chosen to reflect the a-priori assumption of the 50% point being that of 845 msec. This meant that the 25% staircase started at 508 ms (a shorter response would increase duration by 80 ms, a longer response would decrease duration by 240 ms), and the 75% staircase started at 1162 ms (with an increase of 240 ms, and a decrease of 80 ms). Each staircase consisted of 25 oddball presentations and an experimental block ended when all responses were gathered.

##### 2.1.4.1 Transcranial magnetic stimulation

Each session consisted of five phases: a pre-stimulation run of the time perception (TP) task, 10 minutes of rTMS at the first site, a post-stimulation run of the TP task, a 23 minute break, 10 minutes of rTMS at the second site, and a second post-stimulation run of the TP task. During stimulation participants’ heads were stabilized using a chinrest. rTMS was administered using a Magstim Rapid^2^ (Magstim) connected to a Magstim Air-Cooled Double 70mm Coil System. TMS stimulation was a single continuous stream of TMS pulses at 1-Hz. Based on prior work [17] stimulation intensity was set to 65% of total machine output, corresponding to a field strength of 1.9 T for each pulse. The coil was localized to electrode site O1 during occipital stimulation, and electrode site P4 was used to approximate the location of rIPS.

### 2.5 EEG acquisition

EEG was recorded during the entirety of the experimental session. We recorded EEG from 27 scalp locations using Ag/AgCl electrodes (Fp1, Fp2, F7, F3, Fz, F4, F8, Fc5, Fc1, Fc2, Fc6, T3, C3, Cz, C4, T4, Cp5, Cp1, Cp2, Cp6, T5, P3, Pz, P4, T6, O1, and O2 according to the extended 10/20 international system). All electrodes were reference to the right mastoid and CPz was ground. Vertical electrooculogram (VEOG) was recorded with two electrodes placed above and below the right eye. The EEG was amplified and recorded with a full-band DC-EEG system (neuroConn GmbH, Llmenau, Germany) with a sampling rate of 2048 Hz and 300 Hz anti-aliasing filter. Electrode impedences were < 10 kOhm prior to recording.

### 2.5 EEG Preprocessing

EEG data was offline processed using the EEGLAB toolbox for Matlab [18]. Data was resampled to 300 Hz and digitally filtered (0.1-Hz. high pass filter, -3 db per octave; 30 Hz. low pass filter; -6 db per octave). Continuous data was epoched into 1 s. segments (-200 to 800 msec) and baseline corrected (-200 to 0 msec). Epochs with extreme values (+/- 60 µV) were removed from analysis, as were epochs clearly tainted by muscle artifacts or environmental noise. This resulted in the removal of approximately 10% of trials. Independent components were subsequently calculated using the FastICA implementation [19] of Infomax independent component analysis (ICA; [20]). Components representing eye movement artifacts were identified by visual inspection and removed from the data. Finally, because of marked DC drift and post-TMS shifts in mean amplitude data for each participant was normalized. Each ERP thus reflects the individuals’ channel-by-channel z-scored waveform.

### 2.5 EEG Analysis

In order to test if effects are significantly different between conditions we performed a nonparametric cluster-based permutation test on the evoked response between conditions [21], test based on dependent samples F test with the contrast coefficients selected for the required statistical question. We always used 500 randomizations for the monte-carlo simulation to ensure correct confidence intervals for our significance (i.e. smaller than the threshold to take it above an alpha above .05). For the cluster based approach a non-parametric test is conducted by selected features that cluster over our two dimensions spatial (channels) and time. As such for the spatial clusters a neighborhood structure is required, this neighborhood was defined as electrode locations with a maximal distance of 4 cm, resulting in an average of 4.8 neighbors per channel. As some of the hypothesis have *a priori* spatial effects we will sometimes report statistics from a selection of the Occiptial and Parietal sensors only (P4, Pz, P4, O1 and O2). The statistical tests were performed within the Fieldtrip toolbox [22] and custom written scripts.

### 2.2 Results

Three participants were removed from analysis because of excessive noise in the EEG data.

#### 2.2.1 Behavioral Results

We calculated two dependent measures in analysis of behavioral data: the point of subjective equality (PSE) and the just-noticeable difference (JND). The PSE is defined as the 50% point of “longer” vs. “shorter” judgments of the Weibull psychometric curve, equivalent to the stimulus duration at which the participant perceives the oddball stimulus as equal in duration with the standard. The JND was calculated by subtracting the oddball duration observed at the 25th percentile point of the psychometric function from the duration observed at the 75th percentile, garnering a measure of core sensitivity. Psychometric curves were fitted to the data using IGOR 6.2 sigmoid CurveFit function.

As illustrated in Figure 1B, PSE nominally increased following both parietal and occipital stimulation as compared to baseline. This pattern was statistically assessed in a repeated measures analysis of variance (RANOVA) with a factor for stimulation condition (pre-stimulation, post-occipital-stimulation, post-parietal-stimulation), which revealed a marginally reliable effect of stimulation [F (2,34) = 3.025, p = .062, η2=0.151]. A similar analysis of JND revealed no significant effects [F (2,34) = .392, p = .68, η2=0.023].

**Figure 1:**
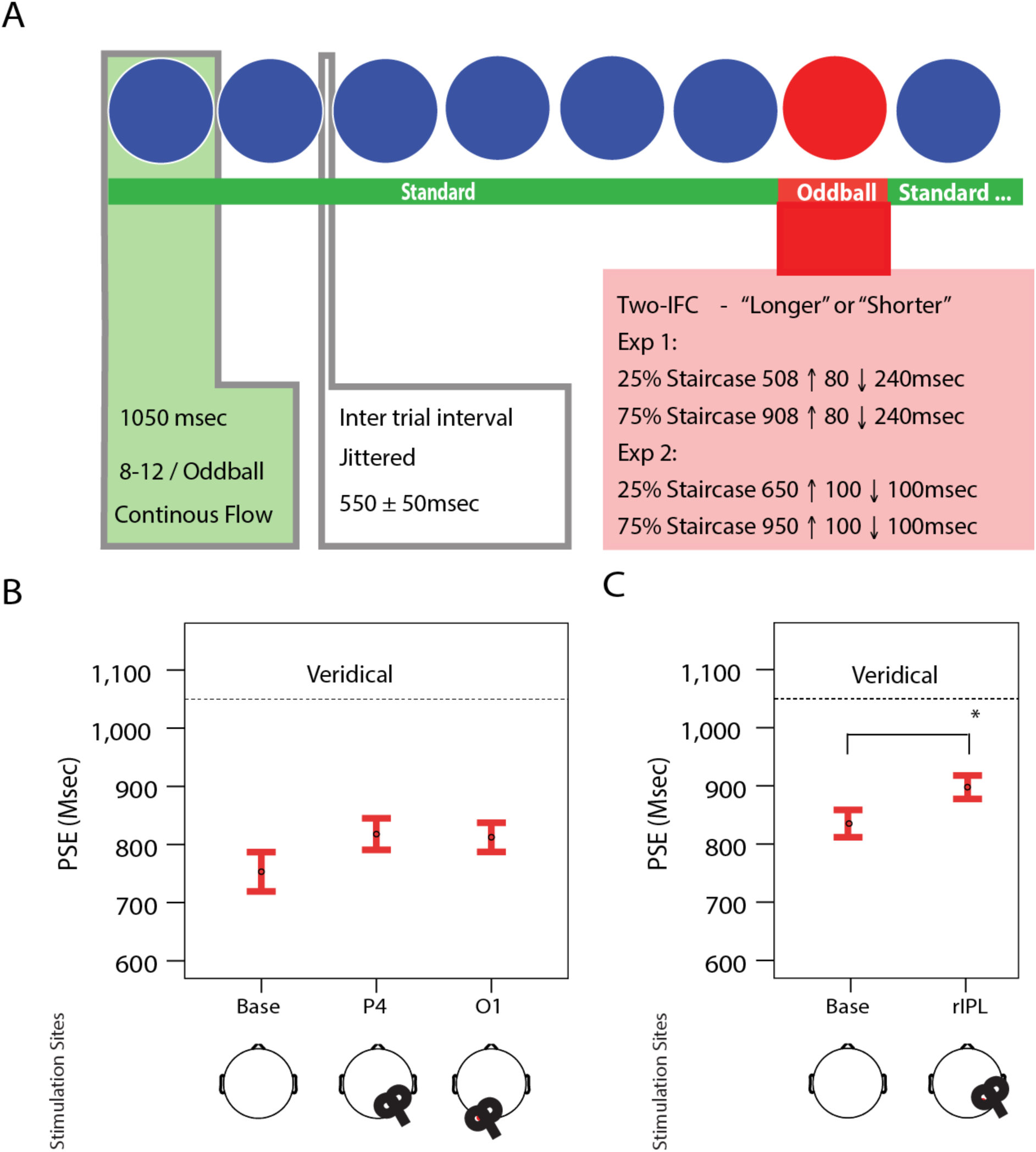
Behavioral Analysis: A: The paradigm of the times subjective expansion. Each trial was wither a standard image that was presented at fixed duration or an oddball image which varied in time based on two staircases that adjusted on response. Participants responded only to the oddball trial, and these responses were to asses if the duration of the oddball was longer or shorter relative to the standard image. B: The PSE effect in Experiment 1. The dashed line corresponds to the veridical duration (i.e. the duration of the standard) and the error bars represent standard error. The below topography show the placement of the TMS coil in the 1Hz pre-task stimulation. C: The PSE effect in Experiment 2. There was a significant difference in the PSE between the baseline and the rIPL stimulation (p = .018) as denoted by the asterix. The dashed line corresponds to the veridical duration (i.e. the duration of the standard) and the error bars represent standard error. The below topography show the placement of the TMS coil in the 1Hz pre-task stimulation.

#### 2.2.2 Electrophysiological Data

Electrophysiological data analysis began with examination of the oddball-elicited ERP as a function of perceived stimulus duration. To statistically assess this pattern, and to test for a possible influence of TMS stimulation site, we conducted a dependent samples F-test across latencies of 300 - 500 ms, with factors for trial type (standard vs. oddball) and stimulation (pre-stimulation, post-parietal-stimulation, post-occipital- stimulation). The cluster-based permutation test revealed one significant clusters for the main effect contrast of duration with frontal and parietal channel significance p = 0.002, corrected (Figure 2 A,B,C). This difference was large and expected. To test for the effect of stimulation we conducted a one-way ANOVA on the oddball and standard separately. For the contrast of stimulation vs. no stimulation no clusters were significant for the oddball: p=0.49 *corrected*; or the standard: p=0.17 *corrected*. Next we examined the contrast for parietal vs occipital stimulation but failed to identify significant clusters in the data. Stimulation thus had no reliable impact on the electrophysiological response to the oddball in this time window.

**Figure 2:**
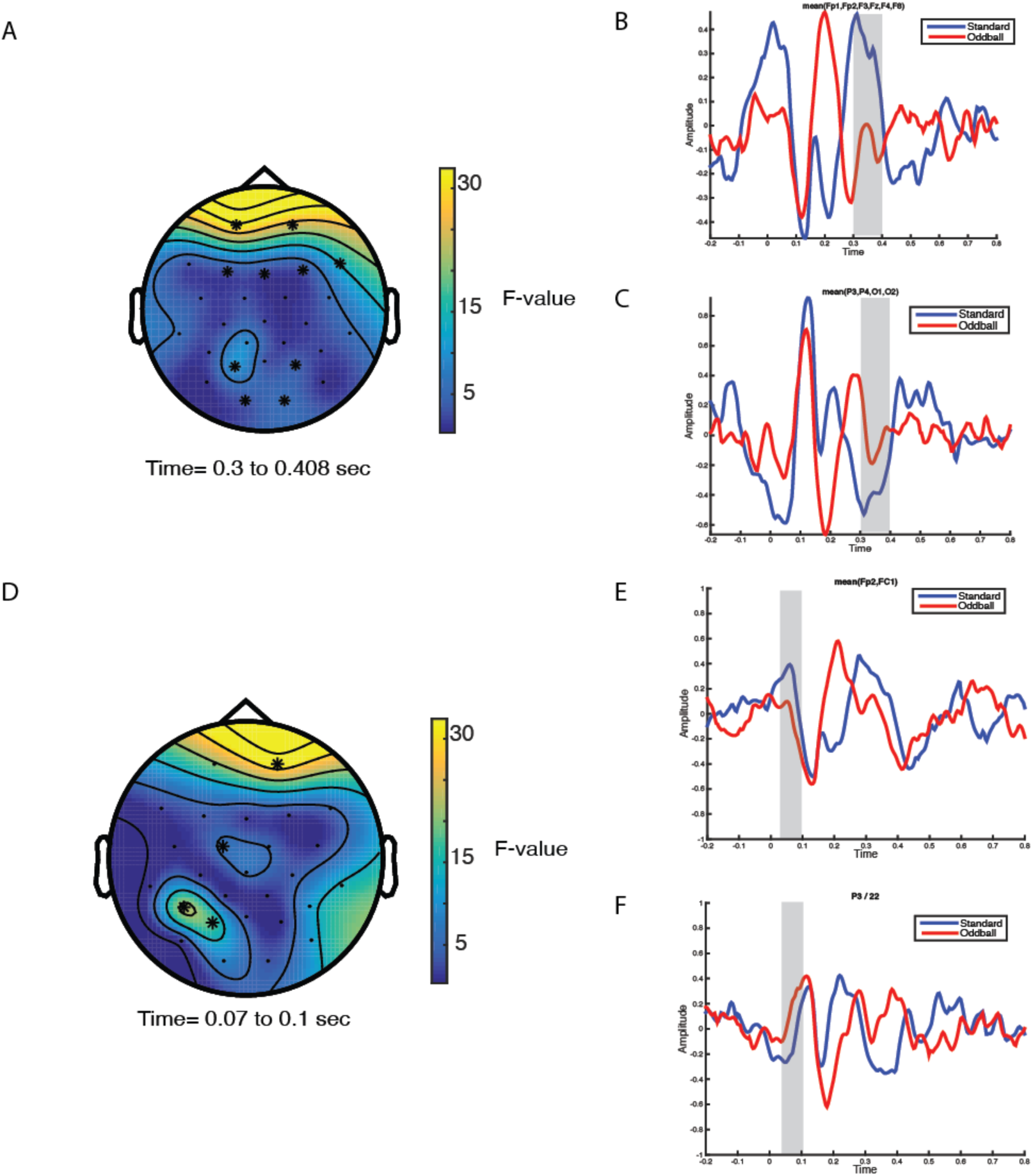
The oddball effect observed in experiment one: A: Topography of the significant effect observed in the 300 to 400 ms window. B: The frontal electrode ERP waveform for the effect seen in A. C: The parietal sensor ERP waveform for the effect seen in A. D: Topography of the significant effect observed in the early 70 to 100 ms time window. E: The effect as seen in the frontal electrode sites. F: The effect at the parietal P3 electrode site.

Due to the hypothesis of repetition suppression [10] a P1 (70 – 130ms) window was also added to this statistical tests, again a main effect of stimulus type was detected with a frontal and parietal electrodes the focus of a divergent effect, p = .004, *corrected* (See Figure 2, D,E,F). All other tests as above showed no significance. The effects in the frontal electrodes seem to be typical of eye movement artifacts, and even though we tried to ensure the removal of these artifacts. Due to this ambiguity and the fact that we did not predict this frontal activity we focus purely on the parietal effects.

After these effects we separated the oddball into longer and shorter, to increase statistical sensitivity we focused on the electrodes that showed significance in the parietal lobe for both the P1 and P3b hypothesis, we therefore selected electrodes P3 P4, O1, and O2. After performing the main effects interactions, and one-way ANOVAs as above, no significant effects were observed.

### 2.3 Discussion

Experiment 1 failed to detect any influence of stimulation on behavioral or ERP measures of perceived oddball duration. However, our results are also inconsistent with the distractor dilation account of the oddball temporal illusion: we find that the amplitude of the oddball-evoked P1 and P3b components predict perceived stimulus duration, thus linking the illusion to variance in target perception.

One possibility is that stimulation in Experiment 1 failed to target the specific aspect of parietal cortex that is involved in time perception and the deployment of attention in time. Our stimulation site was in fact superior to aspects of inferior parietal cortex that have been implicated in time perception in prior research [17,23].

In Experiment 2 we shifted the location of parietal stimulation, targeting a location over IPL inferior to that employed in Experiment 1. The target site was defined as the triangulated center of electrode sites Cp6 P4 and T6. We further adapted our experimental task, adopting a single 1-up 1-down staircase design so as to point staircases at the 50th percentile, rather than the extremes of 25 and 75 percentiles employed in Experiment 1.

## 3: Experiment 2

### 3.1 Methods

All of the methods were the same apart from those outlined below.

#### 3.1.1 Participants

Twelve healthy subjects (7 women) aged 21-31 years (mean 25) with normal or corrected to normal vision participated in the study. All participants gave informed consent. Two were subsequently withdrawn from analysis due to inconsistent results (eg. consistently reporting that an oddball with duration > 1s. greater than the standard was ‘shorter’). Of the remaining 10 participants, 6 were women (19-27 years). The study was approved by the ethical committee of the University of Trento.

#### 2.1.3 Procedure

The experiment was conducted in a single session: Baseline TP Task, 10min rTMS, post-TMS TP Task. The same TP task was repeated each time with the only differences stemming from the adaptive staircases outlined below. The timecourse of the experiment was: TP task (7mins), rTMS Stimulation (10mins), TP task (7mins).

##### 3.1.3.1 Time Perception (TP) Task

The stimuli and procedure were the same as in Experiment 1, however we modified the staircase to a simple 1-up 1-down procedure. The Standard stimuli was presented for 1,050 ms ± 50 ms and throughout the task there was a jittered intra-stimulus-interval of 508 ± 50 ms. Every 8-11 trials the Oddball was presented instead of the Standard. The staircase started at the presumed 50% point of 850ms, and would increase by 100 ms if the participant responded “shorter” and decrease by 100 ms if the participant responded “longer”.

#### 3.4.2 TMS Site Locations

A triangulation of the electrode sites Cp2, P4 and T6.

### 3.2 Results

#### 3.2.1 Behavioral Results

Psychometric curves were fitted to the data using IGOR 6.2 inbuilt sigmoid CurveFit function. A two factor (Baseline, post-rIPL) repeated measures ANOVA was performed on the measures of point of subjective equality (PSE; 50% point of the longer vs. shorter on the psychometric curve) and just noticeable difference (JND; 75% minus 25% on the psychometric curve). There was a significant effect with the PSE measure [F (9,1) = 8.283, p = .018, η2=0.479], and with a smaller mean for Base [*M=839, SE= 25*] than post-rIPL [*M=906, SE= 20*] (See Figure 1). The JND measure showed no significant effect [F (9,1) =1.949, p = .196, η2=0.178].

#### 3.2.2 Electrophysiological Data – ERP Analysis Standard vs. Oddball

Analysis began with examination of the P3b. As in Experiment 1, we calculated mean amplitude for three parietal electrodes (P3, Pz, and P4) across an interval from 300 to 500 ms. A repeated measures ANOVA with factors for stimulus type (Standard, Oddball), stimulation, (baseline, rIPL stimulation), and electrode location ( P3,Pz, and P4) revealed a main effect of stimulus type [F (1,9) =10.024, p = .013, η2=0.556], with oddball stimuli associated with increased positivity, and a main effect of stimulation location [F (1,9) =8.048, p = .022, η2=0.502] but no other effects [Ps < .05]. A similar analysis based on mean amplitude across a 230 – 300 ms revealed no significant effects.

#### 3.2.3 Electrophysiological Data – ERP Analysis Shorter vs. Longer

We additionally analyzed amplitude at parietal sites (P3, Pz, and P4; 300 – 500 ms) as a function of whether participants reported a stimulus to be ‘longer’ or ‘shorter’ than the standards. A repeated measures ANOVA with factors for perceived duration (longer, shorter), stimulation (baseline, rIPL stimulation), for each of the three electrodes separately demonstrated an interaction [P3: F (2,18) =7.060, p = .029, η2=0.469]; Pz: F (2,18) =6.093, p = .039, η2=0.432; P4: [F (2,18) =9.004, p = .017, η2=0.530]. This statistical interaction reflected a pattern wherein oddball-elicited parietal amplitude was smaller when participants reported that the oddball lasted longer than the standard, but only following rIPL stimulation (see Figure 3). A corresponding ANOVA based on mean amplitude from 230 – 300 ms revealed no significant effects, and similar analysis of occipital amplitude (as measured at O1, and O2 from 70 – 130 ms post-stimulus) revealed no significant effects.

**Figure 3:**
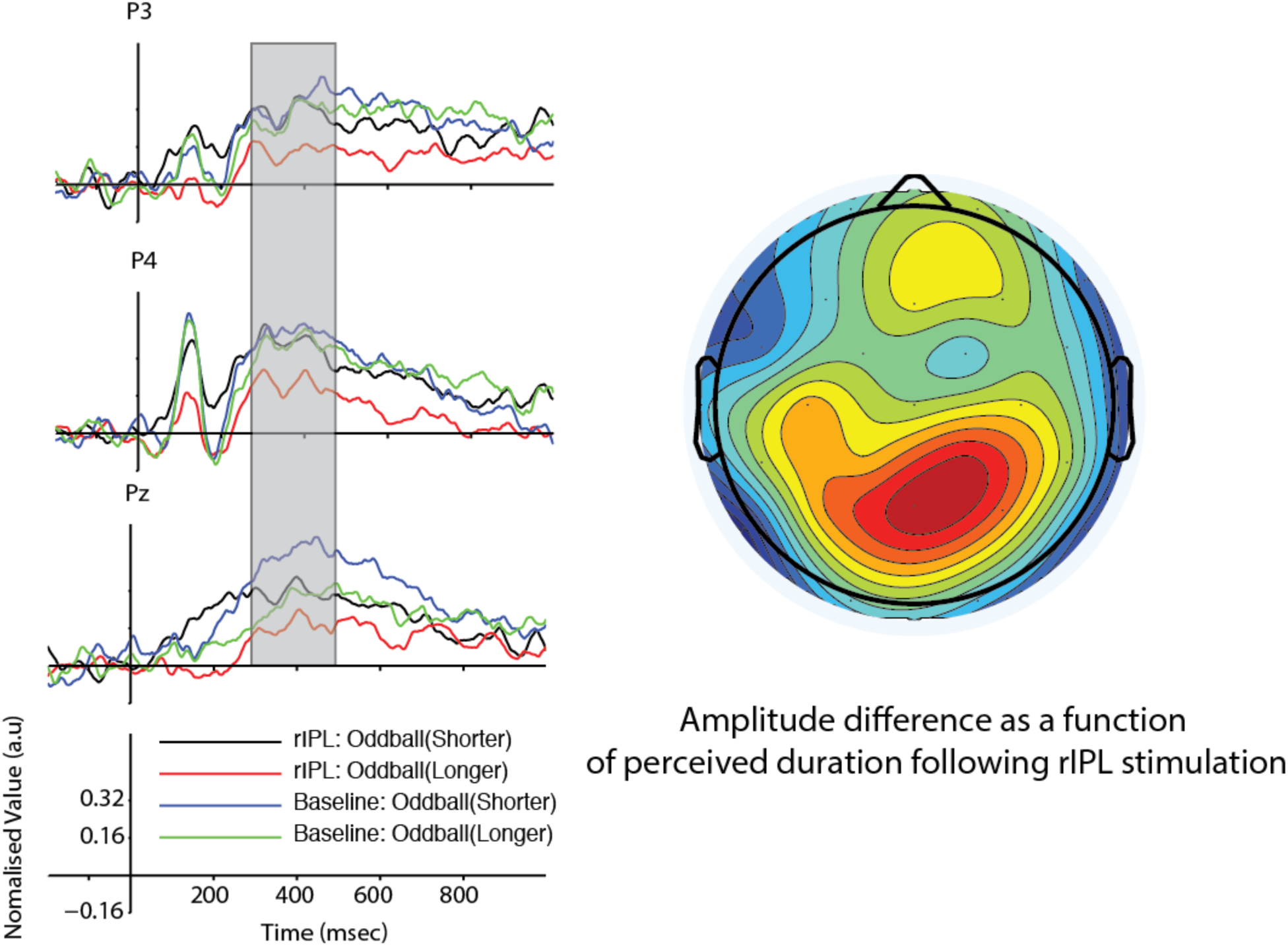
Oddball and standard-elicited parietal ERPs as a function of perceived stimulus duration. Represented in the topographical map is the interaction of perceived duration and stimulation location factors.

## 4: General Discussion

In this study we demonstrated that low frequency rTMS over the right-IPL altered the perception of time in an oddball paradigm. Our results appear selective for stimulation of the rIPL, in that stimulation of the rIPS in Experiment 1 failed to garner a reliable effect in spite of a larger sample size and increased statistical power. This suggests that the rIPL has a discrete role in the perception of time, perhaps particularly when the duration of an event must be judged relative to other events [23]. Inhibitory rTMS of rIPL resulted in a perceived time that was contracted relative to baseline: the oddball had to last 67 ms longer than was the case at baseline to be perceived as equal in length to the standards. Time judgement thus became *more veridical* and the subjective expansion effect of the oddball was reduced.

Results from both Experiments additionally identify for the first time a set of electrophysiological markers of the oddball timing task in normal subjects, highlighting key underlying processes that could influence and dictate the subjective experience of time. The strongest correlate of subjective time expansion in our results is the P3b. The P3b consistently increased in amplitude when participants reported that the oddball lasted less than the standard stimuli. Given that P3b amplitude increases have been linked to stimulus saliency and attentional capture [12], this finding supports the notion that the experience of time is driven in part by the saliency of the oddball and associated capture of attention.

Effects of stimulation emerged only in Experiment 2 when the rIPL was targeted. In the ERP, the primary effect of rIPL stimulation was a decrease in the P3b when the oddball was perceived as being longer than the standard. This suggests a reduction in attentional capture to the oddball, resulting in a more veridical perceptual effect. As this was specific to trials where perceived duration of the oddball was longer, there is also a further suggestion that the P3b effect was stimulus related rather than state related, as a state related distracter effect could lead to an overall reduction of the P3b [24]. Our effects thus appear to be driven by global attentional allocation, consistent with the absence of obvious effects in the ERP prior to the P3b. If effects had emerged prior to the P3b, on the P1 or N2, this would be consistent with the idea that increased stimulus-evoked ‘neural energy’ drives the subjective experience of time, as suggested by Eagleman and Pariyadath (2009). Our null effects on these early components do not rule out the possibility that early evoked activity drives the perception of time, but they do not support it, and the emergence of effects on the P3b without preceding effects strikes us as parsimoniously explained by the ‘attentional capture’ account of time perception.

The right-IPL has been implicated in the processing of temporal information by Battelli and colleagues [25,26] who have proposed that this area is the main locus of the “when” pathway. Our results are consistent with this hemispheric process, and the increased expansion of time. Interestingly, right parietal patients show severe difficulties in a phase discrimination task, a stringent test of relative timing [27]. When asked to discriminate a small counterphase flickering square alternating black and white relative to otherwise identical squares, their phase discrimination threshold was roughly 50% of that of normal subjects (4-5 vs 8 Hz respectively). This reduction of being able to discriminate the onset and offset of the oddball within this task would intuitively lead to an increase in the perceived duration, in agreement with findings that these low-level oscillations maybe coupled to the perceptual windows [28]. In recent experiments Agosta and coworkers [16] provided more compelling evidence of relative time processing within the inferior parietal lobe. The authors tested a large group of right parietal stroke patients on the phase discrimination task [27]. Patients were severely impaired in the timing tasks, while control left parietal patients performed normally. Crucially, the authors tested the direct involvement of the IPL in visual timing by using inhibitory 1-Hz rTMS on healthy subjects. Temporary inhibition of the right IPL (but not of the left IPL) mimicked patients’ performance in normal volunteers. This study directly linked the rIPL to visual timing, confirming the rIPL as a main node in a cortical network specialized in discriminating events across time [25].

Finally, our data support the notion that the processing of time is influenced by the attention capture of the oddball. Indeed, previous studies have implicated the rIPL in attentional allocation. The difficulty though is how to distinguish the temporal mechanism from the attentional mechanism; one way to do this is to see the temporal mechanism as a sub-ordinate to the attentional mechanism. According to this idea, attention can play an inhibitory top-down control on the perception of time, such that disruption of this mechanism through TMS intervention creates a less veridical repetition of time. In this way low-level timing information may be kept “in check” by attention: when attentional control is disrupted, sensitivity to the oddball’s raw salience is altered. The alteration that we observe herein suggests that attention may inflate time in its normal state, but when temporary inhibited with rTMS this exaggeration becomes reduced.

## Conclusion

We find that the effects of times subjective expansion can be influenced causally by rTMS of the rIPL. Stimulation of this location results in a more *veridical* perception of time. This change has an electrophysiological correlate in amplitude of the P3b, a parietal ERP component associated with stimulus salience and attentional capture. These results lead us to conclude that the manipulation of time perception observed in our study reflects a modulation of attentional processing of visual stimuli through stimulation of the IPL.

## Acknowledgments

Lorella Battelli was supported by the Autonomous Province of Trento, Call “Grandi Progetti 2012,” project “Characterizing and improving brain mechanisms of attention – ATTEND”.

